# DNA damage checkpoint dynamics drive cell cycle phase transitions

**DOI:** 10.1101/137307

**Authors:** Hui Xiao Chao, Cere E. Poovey, Ashley A. Privette, Gavin D. Grant, Hui Yan Chao, Jeanette G. Cook, Jeremy E. Purvis

## Abstract

DNA damage checkpoints are cellular mechanisms that protect the integrity of the genome during cell cycle progression. In response to genotoxic stress, these checkpoints halt cell cycle progression until the damage is repaired, allowing cells enough time to recover from damage before resuming normal proliferation. Here, we investigate the temporal dynamics of DNA damage checkpoints in individual proliferating cells by observing cell cycle phase transitions following acute DNA damage. We find that in gap phases (G1 and G2), DNA damage triggers an abrupt halt to cell cycle progression in which the duration of arrest correlates with the severity of damage. However, cells that have already progressed beyond a proposed “commitment point” within a given cell cycle phase readily transition to the next phase, revealing a relaxation of checkpoint stringency during later stages of certain cell cycle phases. In contrast to G1 and G2, cell cycle progression in S phase is significantly less sensitive to DNA damage. Instead of exhibiting a complete halt, we find that increasing DNA damage doses leads to decreased rates of S-phase progression followed by arrest in the subsequent G2. Moreover, these phase-specific differences in DNA damage checkpoint dynamics are associated with corresponding differences in the proportions of irreversibly arrested cells. Thus, the precise timing of DNA damage determines the sensitivity, rate of cell cycle progression, and functional outcomes for damaged cells. These findings should inform our understanding of cell fate decisions after treatment with common cancer therapeutics such as genotoxins or spindle poisons, which often target cells in a specific cell cycle phase.

## INTRODUCTION

The human cell cycle is a complex sequence of molecular events by which cells replicate and segregate their genomic DNA. Cell cycle progression is restrained by multiple checkpoint mechanisms that block transitions between cell cycle phases when cells encounter stressful conditions. For example, DNA damage activates checkpoints that delay cell cycle progression and trigger DNA repair. Under severe stress, DNA damage checkpoints may trigger permanent cellular outcomes such as apoptosis or senescence. Failure to fully activate DNA damage checkpoints can lead to genome instability, as the unrepaired DNA damage can be passed on to the next generation of cells^1–3^. On the other hand, because cells routinely experience low levels of endogenous DNA damage^4,5^, timely checkpoint recovery after DNA damage repair is necessary for continued cell proliferation^6^. Therefore, the balance between cell cycle arrest and recovery must be regulated to continue proliferation in the face of constant exposure to endogenous and exogenous DNA damage sources.

Activation of DNA damage checkpoints is often accompanied by cell cycle arrest, which provides a temporal delay necessary to repair DNA lesions before resuming proliferation^7–10^. The precise timing of events during a DNA damage checkpoint response is not entirely clear. For example, it is not known how abrupt the pause is; whether the pause is a complete halt or a graded reduction in cell cycle progression; or how the duration of the pause changes in response to different DNA damage doses or during different cell cycle phases. Recent live-imaging studies in single cells have shed light on some of the underlying parameters that confer differential sensitivity to endogenous DNA damage. For example, cell-to-cell variation in p21 levels leads to differences in checkpoint stringency (i.e., the robustness of cell cycle arrest in response to DNA damage)^11,12^. However, the underlying parameters that result in differential responses within a cell cycle phase and in response to exogenous DNA damage are generally unknown. Knowledge of these collective dynamical behaviors—which we refer to as DNA damage checkpoint dynamics—is necessary for understanding the relationship between the DNA damage response and functional cellular outcomes such as permanent cell cycle arrest. From a translational perspective, understanding DNA damage checkpoint dynamics could help explain the variability in cellular responses to many chemotherapeutic drugs, which act mainly by inducing DNA damage and interfering with cancer cell proliferation.

Among the various forms of DNA damage, DNA double strand breaks (DSBs) are one of the most harmful types of lesions^13,14^. DSBs can give rise to chromosome rearrangements and deletions that can subsequently lead to cancer. The functional effects of DSBs can vary depending on the cell cycle phase in which the damage was incurred. For example, it has long been known that cells in different cell cycle phases show drastic differences in survival under the same levels of ionizing radiation, which induces DSBs^15^. The underlying basis for these differences in cell survival is not clear. One explanation for the observed differences in survival among the cell cycle phases is that each phase employs unique molecular mechanisms to enforce the checkpoint. Indeed, the concept of distinct DNA damage checkpoint mechanisms at different stages of the cell cycle, such as G1/S, intra-S, and G2/M, has long existed. Precisely how these checkpoint dynamics are executed throughout each cell cycle phase, however, is not clear and requires real-time analysis of DNA damage checkpoint responses. Thus, a quantitative understanding of DNA damage checkpoint dynamics is crucial to linking the molecular pathways to predicted cell cycle outcomes. This knowledge would not only help explain the difference in cell viabilities in response to DNA damage in different cell cycle phases but would greatly inform efforts to predict clinical outcomes of genotoxic chemotherapies, which often target cells in a specific cell cycle phase.

Here, we use fluorescence time-lapse microscopy to reveal the dynamics of DNA damage checkpoints for three major phases—G1, S, and G2—by monitoring cell cycle progression upon acute DNA damage in otherwise unperturbed, asynchronously-dividing single cells. In response to DSBs, each cell cycle phase shows distinct DNA damage checkpoint dynamics. The G1 checkpoint employs a complete halt at high DNA damage levels but is more permissive to DNA damage than the G2 checkpoint because it is temporally located well before the G1/S boundary. The S phase checkpoint is the least sensitive to DNA damage and does not completely halt cell cycle progression, but rather continuously slows the rate of progression throughout the remaining duration of S phase. The most stringent checkpoint occurs during G2, which abruptly and completely halts cell cycle progression by imposing a delay that is linearly correlated with the amount of DNA damage. Because of the internally located checkpoint in G1 and the graded slowdown of cell cycle progression in S phase, both phases show a strong dependence between the timing of DNA damage within the phase and the functional outcome in response to DNA damage. Taken together, our results argue that the timing of DNA damage relative to cell cycle phases determines the sensitivity, dynamics, and functional outcomes of checkpoint responses.

## RESULTS

### A quantitative model of DNA damage checkpoint dynamics and cell cycle phase transitions

DNA damage checkpoint regulation is a dynamic process that needs to adapt to changing environments. These checkpoints must balance the activation and the inactivation of arrest mechanisms in order to prevent lethal consequences of DNA DSBs while allowing cell growth and proliferation. When damage occurs within a cell cycle phase, checkpoint mechanisms halt cell cycle progression until the damage is repaired before allowing transition to the next phase (Figure 1A). To gain a systematic understanding of DNA damage checkpoint dynamics, we developed a quantitative, cell cycle phase-specific checkpoint model that was based directly on observed cell cycle transitions in asynchronously proliferating cell populations. For asynchronously proliferating cells that experience DNA damage at different times during a given cell cycle phase, DNA damage checkpoint dynamics can be visualized by a “transition curve” that captures the rate of transition out of a specific cell cycle phase. To be precise, a transition curve plots the cumulative percentage of cells that transition into the next cell cycle phase against the post-damage transition time—the interval duration between the time of damage and the moment of transition into the next phase (Figure 1B). The plateau in each transition curve represents the time at which nearly all cells have successfully transitioned to the next phase. An advantage of this data-driven model-building approach is that it requires only three conservative assumptions. First, a cell exists in a single, discrete state (cell cycle phase) at a given time^16,17^. Second, the transitions between states are complete and irreversible^18–21^. Third, DNA damage can affect the timing of these transitions through activation of checkpoint mechanisms^22^.

**Figure 1.**
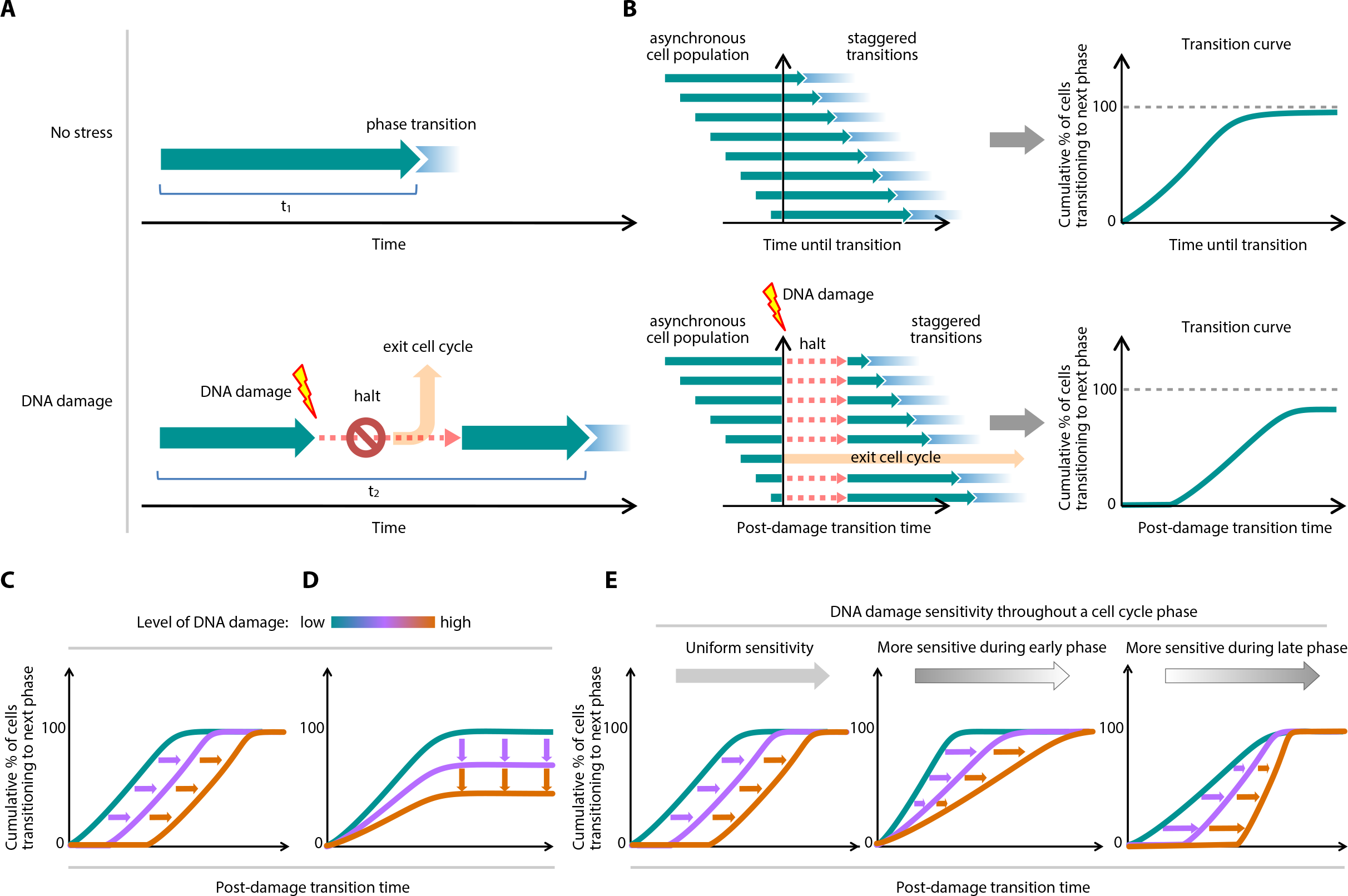
Model of DNA damage checkpoint dynamics. A. “Arrest-and-restart” model of cell cycle checkpoint dynamics in response to DNA damage. The green arrow represents unrestricted progression of a cell cycle phase through time. In the absence of DNA damage, cells progress readily through a phase with duration t_1_. When damage occurs, progression through a phase is temporarily interrupted, resulting in an increase in phase duration, t_2_.
B. Schematic of a transition curve. A transition curve is obtained by measuring the post-damage transition times of an asynchronous population of cells that are damaged at different points during a particular cell cycle phase. Each green arrow represents an individual cell’s progression through a particular cell cycle phase. The transition curve plots the cumulative percentage of cells that have successfully transitioned to the next phase as a function of time.
C. Transition curves expected from a graded halt duration response to DNA damage level. Increasing damage causes a corresponding delay before cells transition to the next cell cycle phase.
D. Transition curves expected from a graded permanent arrest response to DNA damage level. Increasing damage results in fewer cells transitioning to the next phase.
E. Expected transition curves when DNA damage sensitivity varies throughout a cell cycle phase. The intensity of grey arrows indicates the DNA damage sensitivity. Increased sensitivity during the early parts of a phase allows cells that are already near the end of the phase to transition more readily (middle plot). Greater sensitivity in the later part of a phase results in an immediate delay to transition of the population (right plot).

Under this DNA damage checkpoint model, there are two parameters that may vary in response to DNA damage levels. First, the duration of the delay in cell cycle progression can lengthen with increasing amounts of DNA damage. This would correspond to a rightward shift of the transition curve (Figure 1C). Second, when the damage is too excessive to be repaired, a cell may exit the cell cycle and enter permanent cell cycle arrest. As increasing levels of DNA damage increase the number of cells entering permanent arrest, the transition curve plateau lowers (Figure 1D). Under this model, cells can cope with increasing amounts of damage either by lengthening the duration of a temporary arrest, increasing the probability of entering permanent arrest, or both. Importantly, by analyzing an asynchronous population of cells, one can gain insight into the dependency of DNA damage checkpoint sensitivity on the precise time within a cell cycle phase, since the damage is essentially randomly distributed along the time axis of each cell cycle phase. For example, in the case of damage level-dependent delay, as in Figure 1C, the same amount of damage causes a fixed delay duration independent of the timing of damage within the cell cycle phase (Figure 1E, left panel). When in a cell cycle phase whose early part is more sensitive to damage, cells that are damaged during early phase are arrested for longer durations than cells damaged near the end of the phase, resulting in a less steep transition curve. (Figure 1E, middle panel). Similarly, a cell cycle phase that is more sensitive toward the end of the phase shows steeper transition curves (Figure 1E, right panel).

### Quantification of cell cycle progression in individual, asynchronously proliferating cells

To test whether cells employ a dose-dependent arrest duration or a dose-dependent permanent arrest, we developed a fluorescent reporter system that allows real-time monitoring of cell cycle phase transitions in individual proliferating cells. Our goal was to quantify the duration of time a cell spends in G1, S, and G2-M. These three durations were quantified by recording four time points for each cell: the beginning of the cell cycle, the onset of S-phase, the end of S-phase, and the end of the cell cycle (Figure 2A). The beginning and end of each cell cycle was recorded as the time of telophase of the mother cell and the time of telophase of the given cell, respectivly^23^. These two measurements indicate the total cell cycle duration for each cell. To determine the beginning and end of S-phase, we developed a modified reporter for S phase progression by fusing the red fluorescence protein mCherry to the proliferating cell nuclear antigen (PCNA) (PCNA-mCherry; Figure 2B). Multiple studies have shown that PCNA exhibits distinct punctate localization patterns at specific times within S-phase (e.g. sub-phases), and PCNA returns to diffuse nuclear localization upon exit from S phase^24–33^.

**Figure 2.**
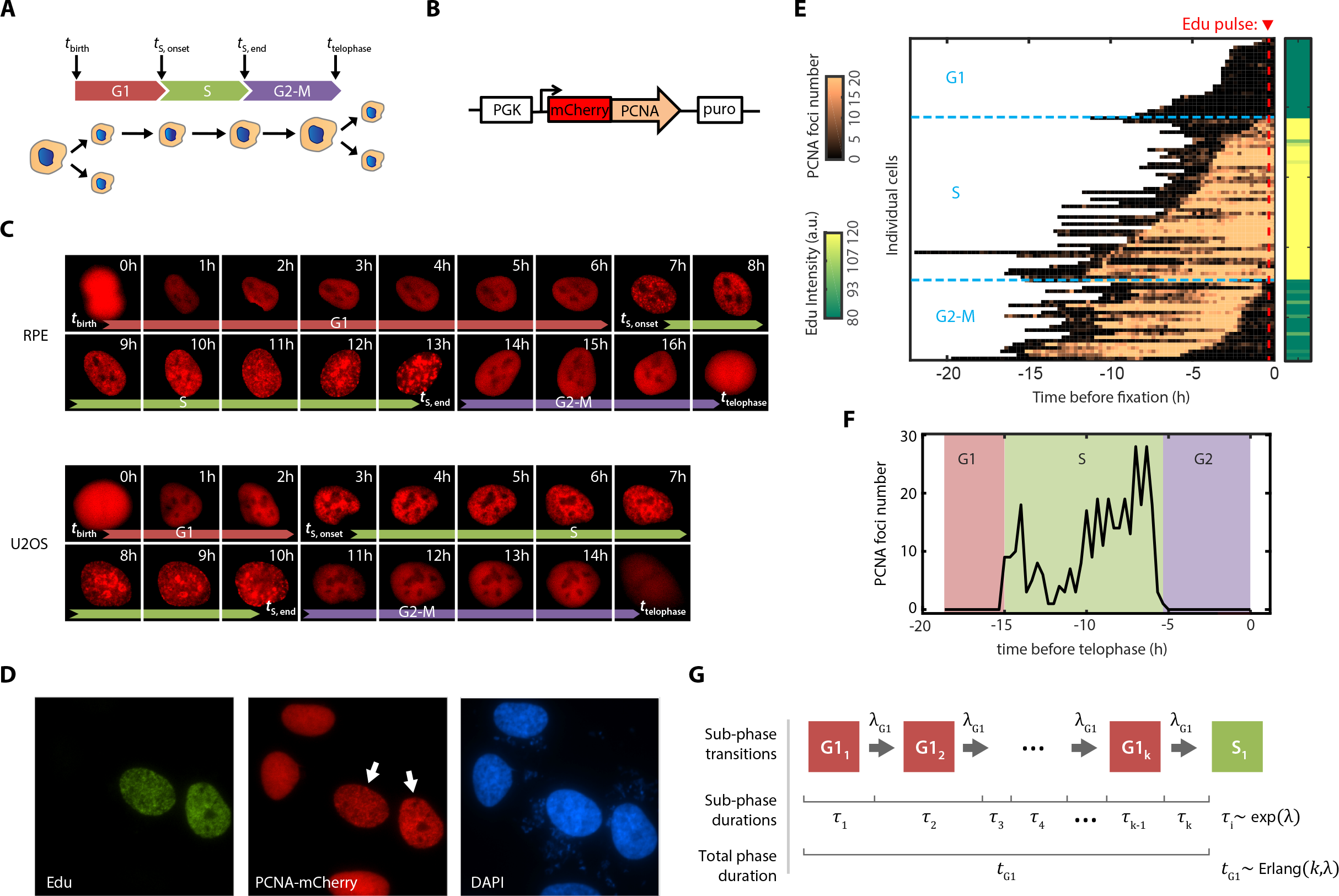
Measuring and modeling cell cycle phase progression in single cells. A. Cell cycle diagram showing three phases—G1, S, and G2-M—defined by four measured time points.
B. Schematic showing the construct design for PCNA-mCherry reporter
C. Sequence of live cell images of RPE and U2OS cells expressing the PCNA-mCherry reporter.
D. Immunofluorescence (IF) staining for EdU incorporation in the reporter cell line. U2OS cells expressing PCNA-mCherry were pulsed with 10 μM EdU for 1 hour, fixed, and stained for EdU and DAPI. White arrows indicate S phase cells with punctate PCNA pattern.
E. Live cell imaging combined with fixed-cell IF staining with EdU incorporation. Asynchronously cycling U2OS cells expressing the PCNA-mCherry reporter were imaged for 24 hours, and the number of PCNA foci in each cell was quantified every 15 mins. At the end of the movie, cells were pulsed with 10 μM EdU for 15 minutes, fixed, and quantified for EdU incorporation.
F. Method for S phase identification based on PCNA-mCherry foci trajectories. The plot shows a typical single-cell time series of PCNA foci number. The onset and the end of S phases were identified by the initial maximal increase and final maximal drop, respectively, in the number of PCNA foci.
G. Schematic of computational model for cell cycle progression. G1, S, and G2-M phases were each modeled as a series of *k* identically distributed subphases with exponential waiting times of rate. λThus, the total duration of each phase followed an Erlang distribution with shape parameter *k* and scale parameter 1/λ. DNA damage was modeled either by interrupting or slowing the rate of subphase progression (see **Supplemental Materials**).

To investigate both functional DNA damage checkpoint dynamics and to understand how disrupted checkpoints can lead to distinct dynamics, as is often found in cancer cells, we stably expressed the PCNA-mCherry reporter in both retinal pigmented epithelial (RPE-1 hTERT, abbreviated RPE) cells and in the osteosarcoma cell line U2OS. RPE cells are non-transformed human epithelial cells immortalized with telomerase reverse transcriptase with intact checkpoints, whereas U2OS cells are a transformed cancer cell line which has an attenuated p53 response and an unstable G1 checkpoint due to the expression of truncated Wip1 and a p16 deficiency, both of which are important for overriding the G1 checkpoint^34–37^. We imaged freely-proliferating cells through multiple cell divisions without affecting the cell cycle distribution properties (**Figure S1A**) or the cell cycle length (**Figure S1B**) as measured through flow cytometry and a traditional cell counting approach, respectively. These controls indicate that time-lapse imaging did not significantly alter cell cycle progression. As expected, we found that PCNA-mCherry formed replicative foci that appeared abruptly at the onset of S-phase and disappeared abruptly at the end of S-phase during the transition to G2 (Figure 2C). The emergence of PCNA-mCherry foci correlated strongly with EdU incorporation, while cells in other phases showed diffuse nuclear fluorescence (Figure 2D-E). This pattern of foci accumulation reliably identified the beginning and end of S-phase with 15-minute precision (Figure 2E-F, **Figure S2**). Thus, we established a system that quantifies, for individual cells, the duration of time spent in G1, S, G2-M, and, thus, the entire cell cycle.

### Computational model of cell cycle progression

We then developed a computational model to stochastically simulate cell cycle progression of RPE and U2OS cells with the goal of eventually understanding how DNA damage affects the rate of progression. We model each cell cycle phase independently as a series of steps, with total number of *k* steps, and a progression rate (λ) through each step (Figure 2G, see **Supplemental Materials**). Therefore, each step of a cell cycle phase can be described as a Poisson process, and the total cell cycle phase duration can be modeled as an Erlang distribution^38^. Because the model is characterized by a sequence of steps with the same rate, it is straightforward to introduce rate changes into cell cycle progression. By fitting the experimentally measured distributions of cell cycle phase durations, we obtained two parameters for each phase: shape (*k*)—which can be interpreted as the number of steps within a cell cycle phase—and scale (1/λ)—which can be interpreted as the average timescale for each of the steps (**Figure S3**, upper panels). Using the estimated parameters, we were able to accurately simulate the cell cycle phase transitions under basal conditions in an asynchronous population with phase durations drawn from the fitted Erlang distribution (**Figure S3**, lower panels).

### Each cell cycle phase displays distinct DNA damage checkpoint dynamics

With this reporter system and computational model in place, we experimentally determined DNA damage checkpoint dynamics for each cell cycle phase in individual RPE and U2OS cells. We induced DSBs in asynchronously dividing cells using the radiomimetic drug neocarzinostatin (NCS) and recorded the cell cycle phase in which the damage occurred. Confirming that NCS induced similar numbers of DSBs in each cell cycle phase (**Figure S4**) without affecting the PCNA-mCherry reporter’s accuracy (**Figure S5**), we then followed each treated cell and measured the time until that cell exited its current cell cycle phase and transitioned to the next cell cycle phase. This analysis produced a transition curve for each dose of NCS and cell cycle phase (Figure 3A-B, solid lines). With no external stress, nearly all of the RPE and U2OS cells transitioned to the next phase (>99%). As the dose of NCS was increased, however, two effects were observed. First, for cells damaged in G1 or G2-M, increasing NCS concentrations reduced the number of cells that completed transitions within the 48-hour observation window, indicating a dose-dependent increase in the permanent arrest probability. Secondly, high DNA damage doses increased the time delay before half of the phase transitions occurred, particularly for cells damaged in G2-M, indicating a dose-dependent increase in delay duration.

**Figure 3.**
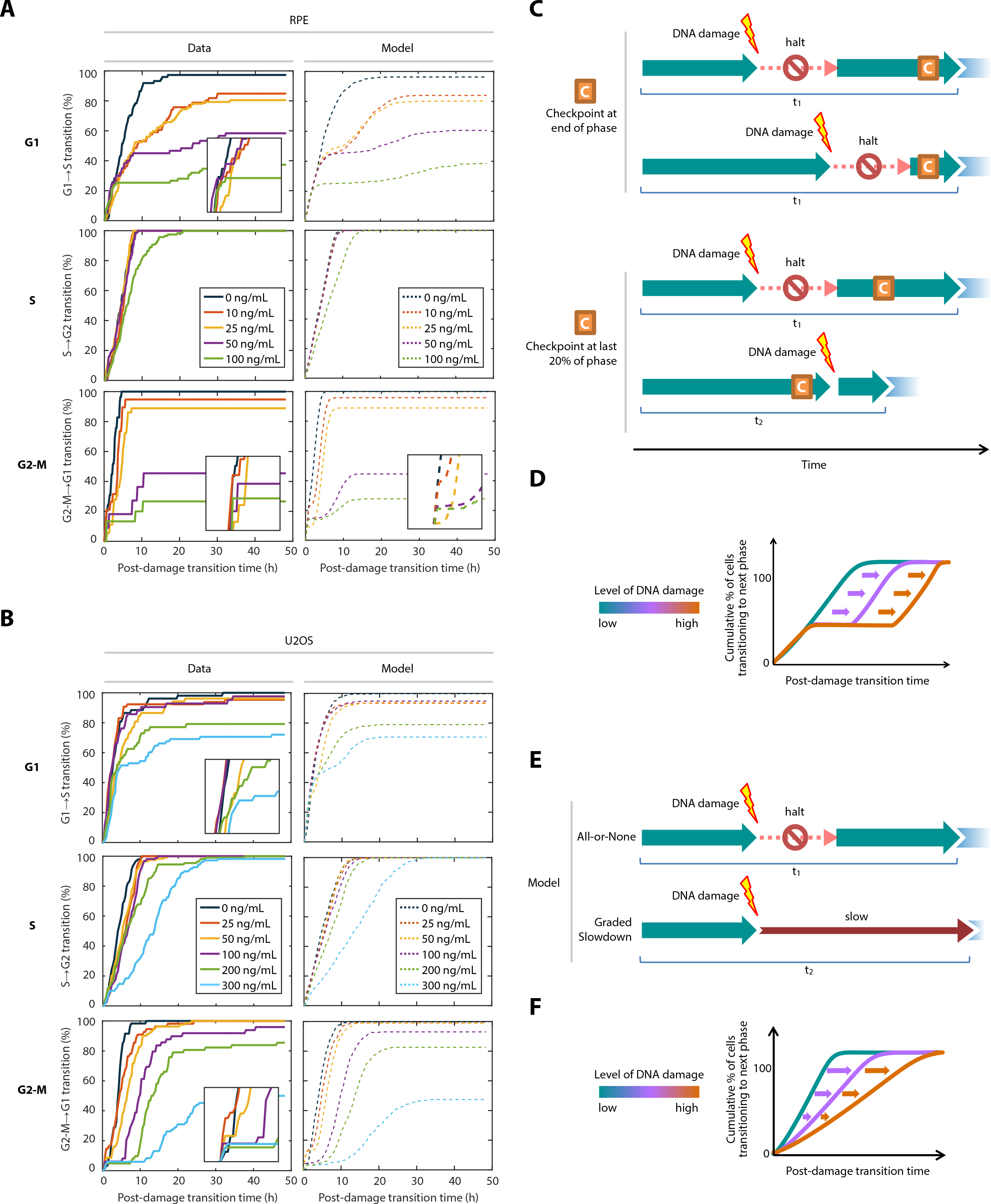
Each cell cycle phase exhibits distinct DNA damage checkpoint dynamics. A-B. Cell cycle phase-specific transition curves in response to acute DNA damage in (A) RPE or (B) U2OS cells. *Left panels and solid lines*: transition curves for asynchronously dividing cells treated with NCS at the indicated concentrations during G1, S, or G2-M. For each cell, we quantified the time interval between NCS treatment and the transition from its current cell cycle phase to the next phase. *Right panels and dashed lines*: Best fit lines of the experimental data to the refined DNA damage checkpoint model that incorporated both the flexible checkpoint location and the possibility of graded slowdown checkpoint kinetics (S phase) described in Figure 3C-F. RPE G1: n=74, 66, 82, 60, 75; S: n= 139, 97, 98, 69, 118; G2-M: n= 29, 19, 36, 11, 15 (order corresponds to 0 ng/mL, 10 ng/mL, 25 ng/mL, 50 ng/mL, 100 ng/mL). U2OS G1: n=52, 65, 52, 42, 48, 68; S: n= 140, 120, 119, 160,109, 109; G2-M: n= 59, 55, 81, 49, 62, 84. (order corresponds to 0 ng/mL, 25 ng/mL, 50 ng/mL, 100 ng/mL, 200 ng/mL, 300 ng/mL). C. Schematic of models for variable DNA damage checkpoint location. The model incorporates a temporally located commitment point. *Upper plot:* End of phase, where the commitment point is located at the end of the phase (i.e., 100% of the way through). *Lower plot:* Last 20%, where the commitment point is located at 80% of the way through the phase. Cells that are damaged after this time point transition to the next phase without arrest. D. Expected transition curves under the model of a DNA damage checkpoint that is internally located. The initial increase in transition percentage for all damage levels represents the fraction of cells that were beyond the commitment point. E. Schematic of models for DNA damage checkpoints with different kinetics. *Upper plot*: A checkpoint that is enforced in an all-or-none manner halts the cell cycle progression completely upon DNA damage. *Lower plot*: A checkpoint that is enforced by slowing down the rate of cell cycle progression upon DNA damage without completely halting the progression. F. Expected transition curves under the graded slowdown kinetic checkpoint model. The change in slopes among different DNA damage levels results from differences in the time of damage for individual cells. Cells damaged earlier in the phase experience a longer delay.

Several additional interesting observations emerged from these transition curves. In both RPE cells, which had a functional G1 checkpoint, and U2OS cells, which have an unstable G1 checkpoint^34,35^, the G1 transition curves exhibited a hyperbolic shape without an immediate delay (Figure 3A-B). Surprisingly, cells damaged during S phase transitioned to G2 without interruption for all but the highest NCS levels, indicating a low sensitivity of the S phase checkpoint to DSBs. The few S-phase transition curves that did show a response to NCS also demonstrated a hyperbolic shape similar to the G1 cells. In addition, almost all RPE and U2OS cells (>99%) damaged during S phase with the highest NCS concentrations were permanently arrested in the subsequent G2-M phase. We observed comparable or even higher arrest probabilities of G2-M arrest after S phase damage than in cells damaged during G2-M with lower NCS concentrations (**Figure S6**). This pattern suggested that cells relied on the checkpoint in G2-M to prevent catastrophic consequences of DNA damage incurred during S phase. In sharp contrast, cells damaged in G2 or M showed a sigmoidal transition curve with an immediate delay. Although the molecular pathways of the DNA damage response in each cell cycle phase employ redundant mechanisms^9^, these results imply that each phase employs distinct DNA checkpoint dynamics. In addition, we found that the transition curves exhibited qualitatively similar features in response to zeocin, a member of the bleomycin antibiotics that induces DNA damage (**Figure S7**), suggesting that the checkpoint responses were general to DNA damage but not specific to a particular DNA damaging agent.

To explain these complex dynamics within our model framework, we fit our experimental data with the “arrest-and-restart” checkpoint model presented in Figure 1A. The model has two adjustable parameters: the duration of the cell cycle halt, and the percentage of cell entering permanent arrest (**Figure S8**, dashed lines, **Table S1** for fitted parameters, see **Supplemental Materials**). Briefly, the halts were modeled by introducing additional times to the cell cycle phase; the permanent arrest fractions were incorporated by randomly selecting a fraction of cells to exit the cell cycle. Although the model reproduced some features of the transition curves, there were significant discrepancies between the model fits and the data, suggesting that important features of the DNA damage checkpoints were not captured by the “arrest-and-restart” checkpoint model outlined in Figure 1A. For example, cells damaged in G1 transitioned to S phase at the same rate as unstressed cells until ~3 hours after damage, at which point the transition curve began to flatten. This initial rise in transition curves was also observed in G2-M for the first ~1 hour after damage. This behavior could be explained by a checkpoint commitment point that was temporally located roughly 3 hr (G1) or approximately 1 hrs (G2-M) before the end of the phase. Cells that have passed such a DNA damage “commitment point” are already irreversibly committed to transitioning at the time of damage despite experiencing DNA damage. A second feature of the data that was not well captured by the model was the hyperbolic shape of the curves observed in S phase, which often showed gradual rightward shifts at the 50% transition point. These features could be explained by a slowing down—rather than a complete halt—of cell cycle progression when the checkpoint was activated by DNA damage.

### A refined model of DNA damage checkpoint regulation of cell cycle progression

In gap phases, DNA damage checkpoints prevent cells with significant DNA damage from entering the next cell cycle phase. The simplest description of these checkpoints is that they block transition to the next phase regardless of when the damage occurs within the phase (Figure 1A). However, the observed transition curves prompted us to consider the possibility of a “commitment point” located at some internal point of a cell cycle phase after which DNA damage does not prevent a successful transition. The exact temporal location of such a commitment point is unclear. For example, a commitment point could exist at the phase-phase boundaries, resulting in arrest essentially regardless of when the damage occurs (Figure 3C, upper plot). Alternatively, the commitment point may be located somewhere internal to the cell cycle phase (Figure 3C). In the latter case, once a cell passes this point, it would not be subject to arrest by a DNA damage checkpoint in that phase but would presumably affect progression through the next phase. This model would be expected to lead to a delay in the divergence of the transition curves at all DNA damage doses (Figure 3D). Moreover, we also allow that the cellular consequence of the checkpoint activation may not lead to a complete halt of cell cycle progression, which we refer to as “all-or-none” checkpoint activation (Figure 3E, upper axis). Instead, the checkpoint enforcement could induce a partial slowdown of cell cycle progression that adjusts the *rate* of progression relative to the extent of DNA damage level. We refer to this model of checkpoint enforcement as having “graded slowdown” kinetics (Figure 3E, lower axis). Graded slowdown checkpoint kinetics would lead to a decreased slope in the transition curves (Figure 3F).

To distinguish among these possibilities, we sought evidence of an internally located commitment point as well as checkpoint enforcement through graded slowdown kinetics. Computational simulations revealed a qualitative difference in the transition dynamics under different DNA damage checkpoint models (**Figure S9, Supplemental Materials**). For checkpoints (e.g. G2-M) that triggered an abrupt and complete halt, the transition curves exhibited a sigmoidal shape with an immediate delay in transition right after damage (**Figure S9A**). In contrast, under the graded slowdown model, where cells partially slowed down their progression in response to DNA damage, the transition curves were hyperbolic (**Figure S9B**), as seen mostly in S phase and occasionally in G1. Furthermore, simulations that incorporated the internal checkpoint location revealed a delay in divergence of the curves under both checkpoint models (**Figure S9C-D**, comparing red line to other lines) as seen, for example, in the G1→S transition for both RPE and U2OS cells (Figure 3A-B).

We then fit each phase of our experimental data to the refined model of two different kinetics—all-or-none and graded slowdown—and determined the kinetics that gave a better fit. (Figure 3A-B, dashed lines, **Figure S10, Table S2-3**, see **Supplemental Materials**). For G1 in RPE and G2-M in both cell types, the data were consistent with all-or-none checkpoint kinetics. In addition, both G1 and G2-M showed evidence of internally located commitment points in both cell types. We estimated the commitment point location to be ~60% (average among individual estimates from non-zero NCS concentrations, 51%-75%) of the full G1 duration for RPE and ~50% (35%-68%) for U2OS cells. For the combined G2-M phases, we detected commitment points at ~80% (76%-90%) and ~90% (85%-96%) of the full G2-M duration for RPE and U2OS, respectively (**Table S2-3**). Since entry into M-phase is known to temporarily silence the DNA damage response (DDR)^39–44^, this temporal position likely corresponds to the G2/M boundary, which was not precisely resolved in our system. For S phase cells, however, the graded slowdown checkpoint kinetics produced a better fit for both RPE and U2OS, with the commitment point located at 97% (96%-97%) and 96% (90%-100%) of S phase, respectively. Overall, the G1 commitment point estimates showed the highest variability among different NCS concentrations, which could be due to the heterogeneity in G1 durations (**Figure S3**). However, commitment point estimates were in agreement within G1 but significantly different from those of G2. In addition, when we repeated the fitting, the estimates under the highest NCS concentrations fell into a defined distribution (**Figure S11**). Altogether, our results suggested that cells in G1 under high damage levels and all cells in G2-M implemented an all-or-none checkpoint activation whereas checkpoint responses in S phase showed graded slowdown kinetics. Furthermore, the G1 commitment point was located well before the G1/S boundary. In addition, the refined model predicted that both the halt duration and the permanent arrest probability in gap phases increase with the level of DNA damage (**Figure S12**). Strikingly, cells damaged in G1 and G2-M that transitioned into the next phase were significantly affected in the subsequent gap phase, either by prolonging the phase or by increasing the permanent arrest probability (**Figure S13**). These results further suggest that the commitment points in G1 and G2-M both represent a “point-of-no-return” regardless of the levels of DNA damage.

### G1 and G2 checkpoints abruptly and completely halt cell cycle progression

We next sought to verify the predictions of our refined DNA damage checkpoint model. Specifically, the model predicts a dose-dependent halt duration in phases that display all-or-none checkpoint activation such as G1 or G2-M. It also predicts that these checkpoints are enforced at internally located commitment points. This would presumably lead to two populations of cells depending on the timing of DNA damage: cells damaged before the commitment point should be equally delayed in phase progression regardless of when the damage occurs, whereas cells damaged after the commitment point should proceed to the next phase without delay. In contrast, the model predicts that cells damaged in S phase should partially delay their transition to G2 in proportion to the amount of damage. We examined the distributions of post-damage transition times in the three cell cycle phases, which showed evidence for two distinct checkpoint kinetics (Figure 4). Indeed, in the phases where our models predicted an all-or-none checkpoint (G1 or G2), we also observed distinct gaps in post-damage transition times (Figure 4A, 4C). The gap was most dramatic in G2-M, suggesting a checkpoint that imposed a stringent time delay in response to DNA damage; all cells that had not passed the commitment point were arrested for a substantial time. In contrast, cells damaged during S phase showed continuous distributions of post-damage transition times without any obvious gaps, consistent with the model’s prediction of graded slowdown checkpoint dynamics (Figure 4B). We thus concluded that our refined model captured many of the DNA damage response dynamics displayed by proliferating cells.

**Figure 4.**
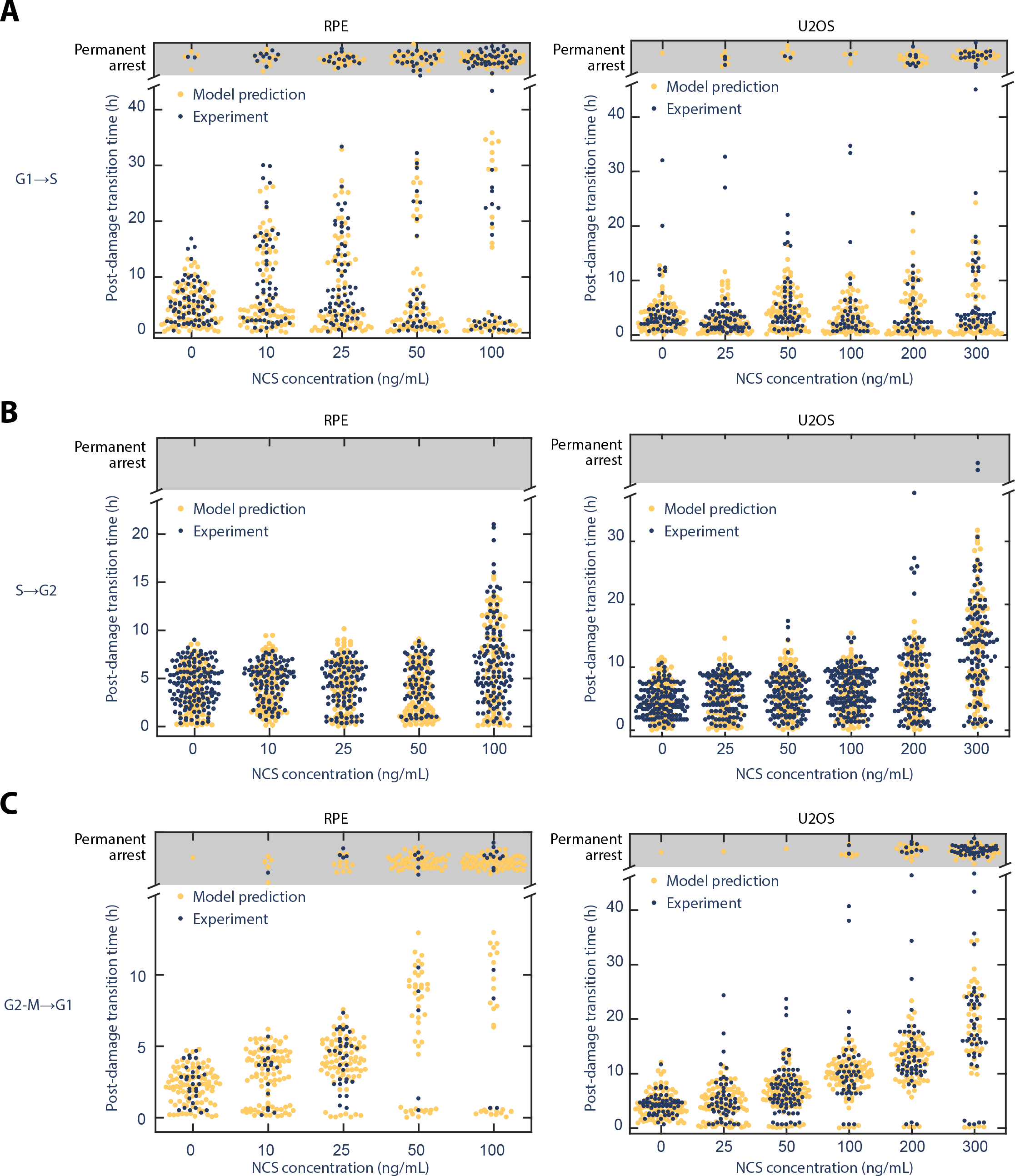
The G1 and G2 DNA damage checkpoints enforce a complete halt to cell cycle progression by an interval that is proportional to the damage level. A. The post-damage G1→S transition time durations of cells damaged anytime during G1 in RPE (left panel) and U2OS (right panel) cells.
B. The post-damage S→G2 transition time durations of cells damaged anytime during S in RPE (left panel) and U2OS (right panel) cells.
C. The post-damage G2-M→G1 transition time durations of cells damaged anytime during G2 or M in RPE (left panel) and U2OS (right panel) cells.

We next characterized the checkpoint kinetics with respect to the arrest durations. Because the G2-M checkpoint provided the strongest evidence of a stringent, complete halt with a late and consistent commitment point, we focused our studies on the G2-M arrest kinetics.

The model predicted that the commitment point locations for G2-M were relatively close to the end of the combined G2-M phase (80% for RPE and 90% for U2OS). Consistent with the model prediction, we noticed only a small population (19% for RPE; 6% for U2OS) of G2-M cells that escaped the checkpoint and divided immediately after the highest two levels of NCS treatment (Figure 4C). These sub-populations that divided soon after DNA damage likely represent cells that had already entered mitosis, since the entry into mitosis has been shown to temporarily inactivate the DDR^9,41–43,45^. Indeed, based on the average G2-M durations for RPE (3.9 hrs) and U2OS (6.5 hrs) (**Figure S3**), the model predicted the checkpoint location to be ~0.7-0.8 hr before division. Since the average M phase lasted approximately 40 mins under basal conditions based on nuclear envelope breakdown revealed by differential interference contrast images (**Figure S14B**), the predicted commitment point location coincided with the G2/M boundary. Although the duration of mitosis in U2OS cells appeared to increase under the highest NCS concentrations (**Figure S14C**), this does not affect the estimation of the commitment point location because that estimation relies on immediately transitioning cells that do not show arrest or delay. Because the DNA damage checkpoint response was presumably activated before reaching the G2/M boundary, we infer that the checkpoint delays observed in our data were enforced entirely within G2 and not M phase. We therefore refer to this checkpoint as simply the G2 checkpoint henceforth.

The G2 checkpoint may impose a constant time delay upon a threshold level of DNA damage, or it may be programmed to calibrate the delay durations according to DNA damage levels, as suggested by the model fitting (**Figure S12A-B**). To distinguish between these possibilities, we examined the time gap between the onset of acute DNA damage and the next transition time point. We found the time delay to be prolonged with increasing NCS concentration (Figure 4C), in linear proportion to the level of DSB incurred (**Figure S15**). Furthermore, the arrest durations were independent of when the cells were damaged within G2 (**Figure S16**). Together, these results suggested that the majority of G2 phase was under the protection of the DNA damage checkpoint. In response to acute DNA damage, the G2 checkpoint abruptly imposed a time delay that was proportional to the level of damage and independent of the progression stage of G2 phase.

### The graded slowdown and non-terminal commitment point location link cell cycle outcome to cell cycle stage

Our findings thus far indicate that each cell cycle phase has distinct DNA damage checkpoint kinetics and commitment locations. Although the two mechanisms are distinct, both the internal checkpoint location and the graded slowdown models predict that an individual cell’s transition kinetics depend on the precise time that the cell experiences DNA damage. In the case of an internally located commitment point, DNA damage leads to arrest only when it occurs before the commitment point. In the case of a graded slowdown mechanism, cells that experience damage earlier during a phase would be expected to take longer to complete that phase since there is more remaining time subjected to the slowdown. We sought to test these predictions with further analyses of cells damaged at different times during each phase. We used asynchronously dividing cells damaged at different points during G1, S, and G2-M phases and plotted the total phase duration against the time already spent in that phase (Figure 5A).

**Figure 5.**
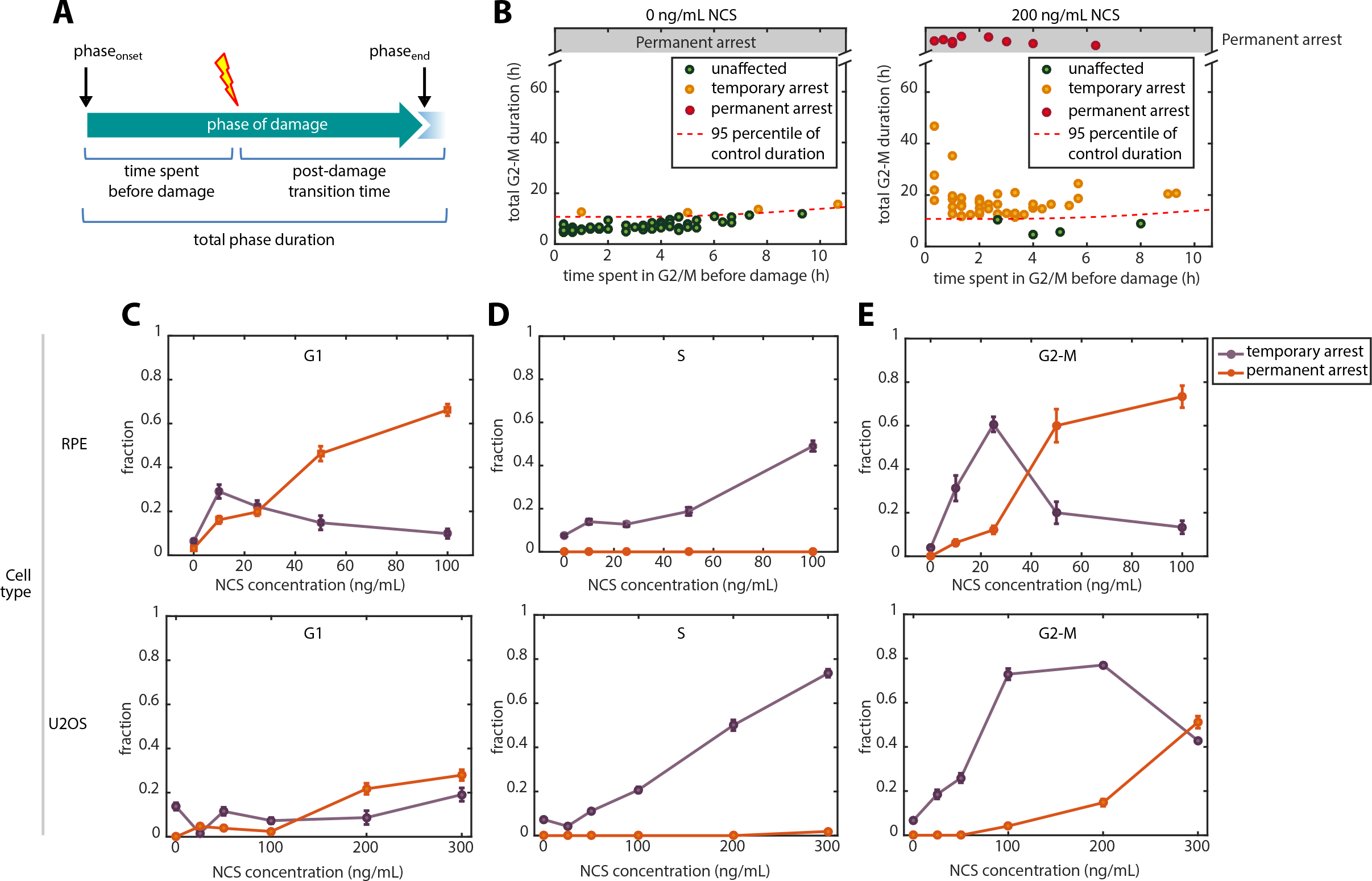
Cell cycle outcomes as a function of cell cycle phase at the time of DNA damage. A.Schematic showing the definitions of time spent before damage, post-damage transition time, and total phase duration. B.Classifying individual cells into three cell cycle outcomes based on phase duration. Here, we show U2OS cells treated during G2-M to demonstrate the classification strategy. Untreated cells (*left*) were used to identify a subpopulation of cells whose phase durations fell above the 95% percentile. These cells were considered to be arrested. (*right*) U2OS cells treated with 100 ng/mL NCS during G2-M showing a large fraction of cells above the threshold identified in the control population. C-E. Phase-specific cell cycle outcomes in response to increasing doses of NCS treatment. Error bars represents binomial standard error. *Upper panels*: RPE. *Lower panels*: U2OS

To investigate the functional outcomes of these checkpoint responses, we categorized cells into three groups: unaffected (cells with total duration not significantly different from the control distribution), temporary arrest (cells with prolonged cell cycle phase but eventually transitioned by the end of the 48-hr experiment, which can be due to either all-or-none or graded slowdown checkpoint kinetics), and permanent arrest (cells that did not transition within 48 hrs) (Figure 5B, **Figure S17**). Although transitions could potentially occur after 48 hrs, most transitions took place during the first 35 hrs with a small number of additional transitions afterwards (Figure 4), indicating statistically different responses between the transitioned and un-transitioned groups.

To assess cell cycle outcomes from DNA damage in different cell cycle phases, we plotted the three categories of cell cycle outcomes as a dose response for each cell cycle phase (Figure 5C-E, **Figure S18**). Several interesting features emerged from this analysis. First, in all three phases, a smaller fraction of U2OS cells were affected by the same NCS concentrations compared to RPE, indicating a higher tolerance to DSBs as expected. Second, S phase cells had the highest NCS tolerance and responded only by temporary slowdown (Figure 5D) but presumably underwent permanent arrest in the subsequent G2 (**Figure S6**). Third, in both G1 and G2, cells preferred temporary arrest response to low NCS levels, but changed to permanent arrest response to high NCS. Unlike during G1, when temporary arrest was never a predominant outcome (always <40%) and the change from temporary to permanent arrest was gradual (Figure 5C), G2 cells preferred temporary arrest in response to low NCS, but displayed a sharp switch to permanent arrest at around 40 ng/mL NCS in RPE and around 300 ng/mL in U2OS. The total fraction of affected cells (cells undergoing temporary or permanent arrest) was higher in G2 compared to G1 in both RPE and U2OS. Thus, the G2 checkpoint is overall the most stringent checkpoint, because fewer damaged cells escaped the checkpoint response. In sum, the results suggested G1 and S phase checkpoints involve a gradual transition to permanent and temporary arrests, respectively, whereas the G2 checkpoint includes a sharp threshold above which cells switch to permanent arrest.

Finally, we asked whether cell cycle outcome depended on the timing of DNA damage within a cell cycle phase. Simulations revealed that, under the all-or-none model, the affected fraction was independent of cell cycle phase stage at the time of damage, as long as the damage occurred before the commitment point (Figure 6A, **Figure S19**, middle panels). In contrast, under the graded slowdown checkpoint model, the fraction of cells affected was inversely proportional to the progress into that phase at the time of damaged (Figure 6A, **Figure S19**, right panels). Consistent with the model prediction, the total S phase durations were linearly and inversely correlated with the time spent in S phase before damage, with a slope consistent with the model predicted slowdown factors (Figure 6B, **Figure S20**). This relationship was in contrast to the all-or-none checkpoint kinetics in G2-M, where the total phase duration was independent of the timing of damage (**Figure S16**). This correlation also argued against the possibility that the decreased slope in S phase transition curves was due to the increased variations in halt duration with increased DNA damage level.

**Figure 6.**
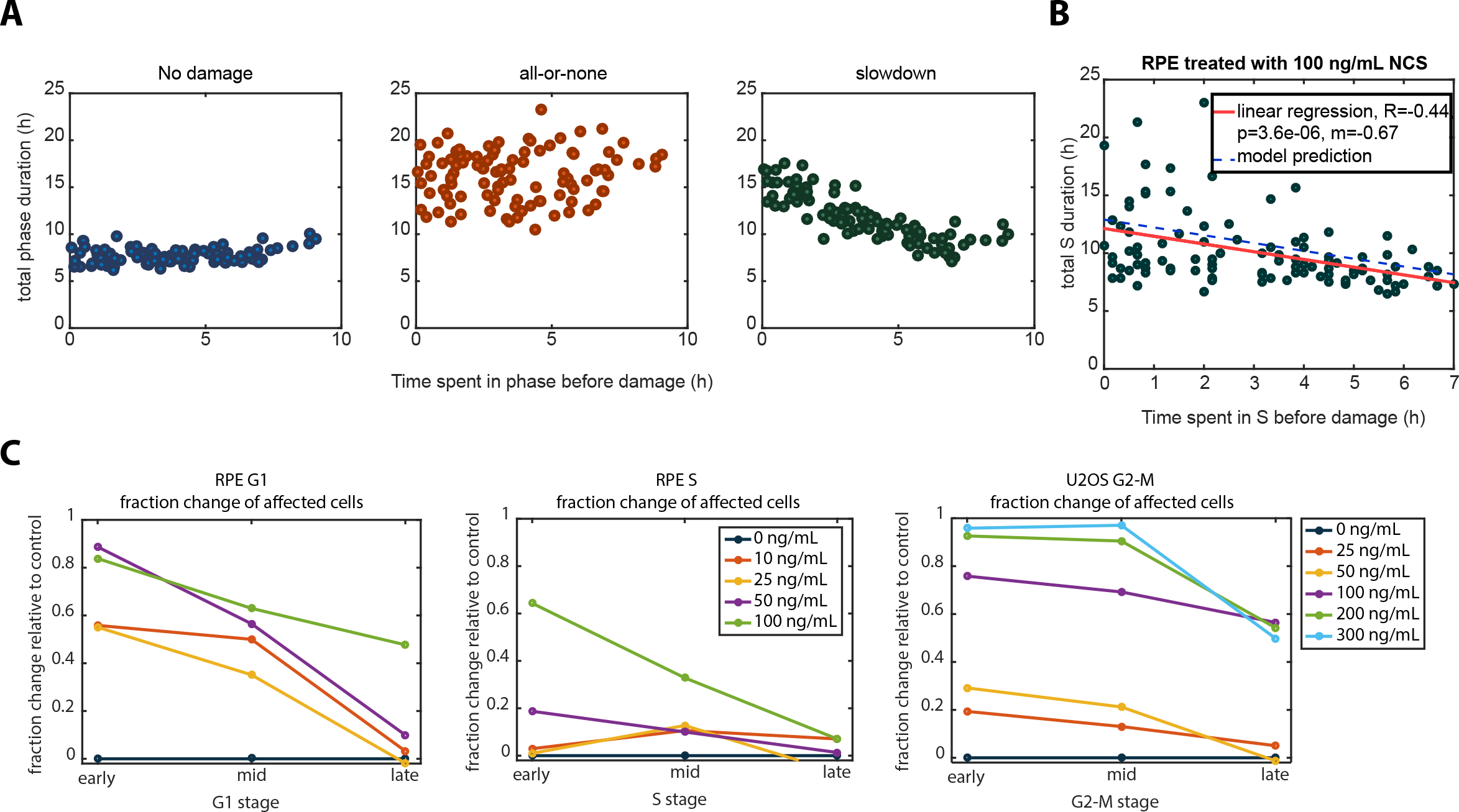
Cell cycle outcomes depend on the timing of DNA damage both within and between cell cycle phases. A. Simulations of cell cycle phase durations as a function of time spent in that phase before damage. Simulations were performed using the fitted parameters (the shape, *k*, and scale, 1/λ) from RPE’s G1 phase distributions without taking into account the commitment point. In the cases with all-or-none checkpoint kinetics, the delay duration was 8 hrs. In the cases with graded slowdown checkpoint kinetics, the slowdown factor was 0.5.
B. The delay in S phase completion linearly depends on the timing of DNA damage within S phase. Experimental data of the total S phase durations as a function of time spent in S phase before damage in RPE cells under 100 ng/mL NCS. Red line represents linear regression of the experimental data. Blue dash line represents the model predictions of the mean S phase durations based on the graded slowdown checkpoint kinetics. The model parameters were based on the RPE’s average S phase duration of 7.7 hrs under basal condition and estimated slowdown factor of 0.60 under 100 ng/mL NCS.
C. Dependency of cell cycle outcome on the stage of cell cycle phase progression. Representative cell cycle outcomes for each phase’s checkpoint are plotted as a function of cell cycle phase stage—early, mid, or late stages. Fraction change relative to control was obtained by calculating the difference between the fraction of cells being affected (either temporary or permanent arrest) under NCS treatment and that without NCS treatment. G1 and S phases are shown for RPE cells, which display more physiological checkpoint responses than U2OS cells. G2 phase is shown for U2OS cells because the RPE G2 duration was too short to be readily subdivided into three stages. Cells were assigned to early, mid, or late stages based on whether the time they spent in that phase fell within the first, second, or third equal partition (tercile) of the theoretical distributions of time spent in each phase, respectively.

When we examined the cell cycle outcome’s dependency on three stages within a given cell cycle phase at time of damage (early, middle, and late), we noted consistent patterns: Cells damaged earlier during a phase were more likely to be affected than cells damaged later (Figure 6C, **Figure S21**). The S phase checkpoint, which utilizes a graded slowdown checkpoint model, showed a linear decreasing effect as a function of time at the highest NCS concentration in RPE (Figure 6C, middle panel). In contrast, G2-M’s early and middle stages, which presumably included only G2 cells, did not show a difference in arrest fraction, which was consistent with the abrupt all-or-none model (Figure 6C, right panel). The decreased fraction in the late stage was mostly due to the insensitivity to DSBs during mitosis. Therefore, this biphasic response in G2-M was characteristic of an all-or-none checkpoint model with a late commitment point. Importantly, G1 checkpoint outcome was inversely dependent on the stage progression (Figure 6C, left panel), consistent with a commitment point substantially earlier than the G1/S boundary.

## DISCUSSION

Here, we used time-lapse fluorescence microscopy to study the dynamics of DNA damage checkpoints in asynchronously proliferating single human cells. We used single-cell data to build a phenomenological model of cell cycle progression that incorporates the time-dependent effects of DNA damage. Our analysis reveals new aspects of cell cycle phase-specific DNA damage checkpoint dynamics (Table 1). Individual phases varied in several features: abruptness of the arrest response (all-or-none versus graded slowdown kinetics), the temporal location of an internally located commitment point, and the types of functional responses (e.g., temporary versus permanent arrest). These results provide a systematic characterization of DNA damage checkpoint dynamics that enhances our current understanding of cell cycle checkpoint kinetics.

**Table 1.**
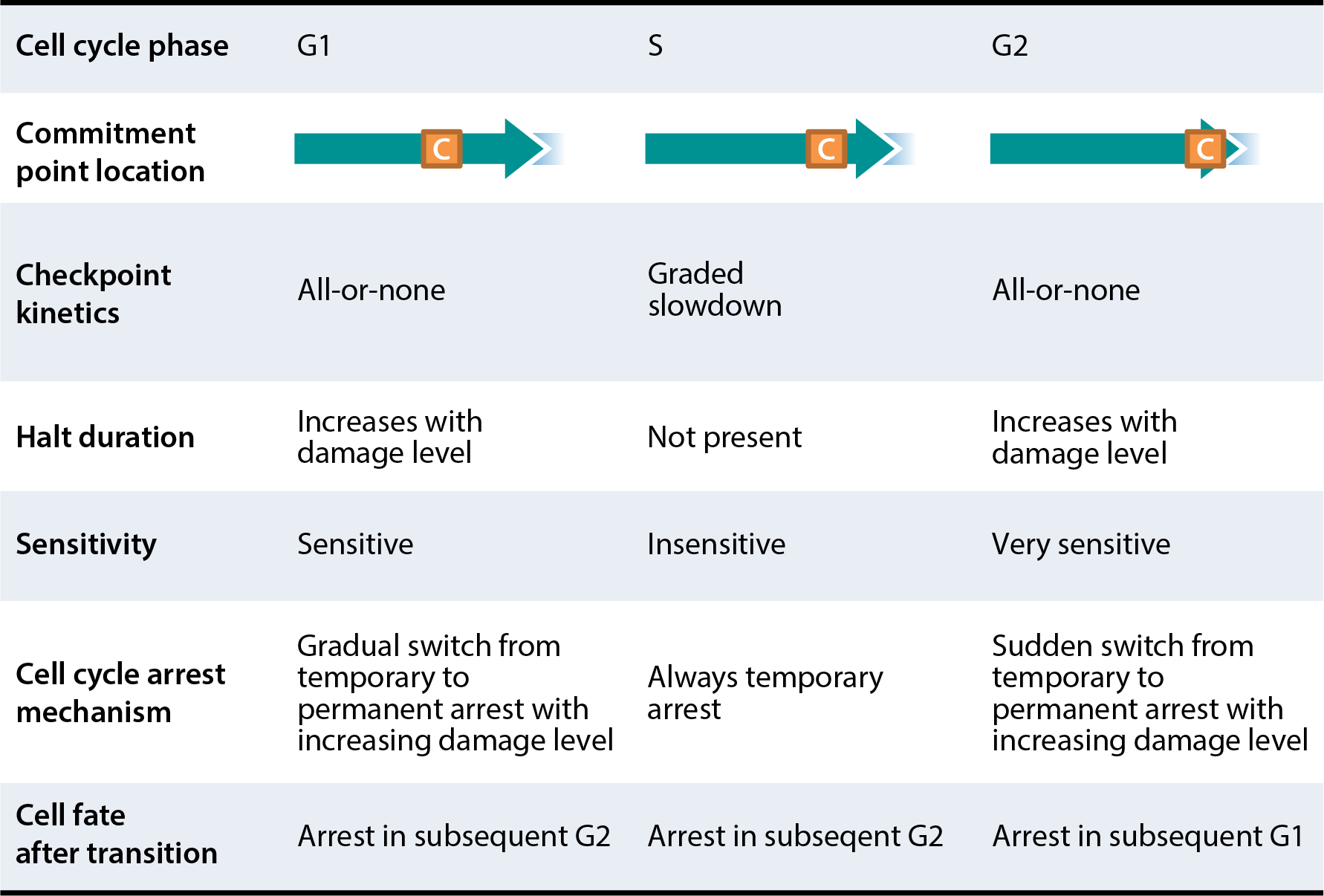
Summary of checkpoint dynamics as a function of cell cycle phase in RPE cells.

The different checkpoint kinetics predicted by our model can be explained by the underlying biological mechanisms of each checkpoint. The transition from G1 to S phase is marked by the activation of DNA synthesis from multiple origins of DNA replication, an event that requires active Cyclin E/CDK2 and/or Cyclin A/CDK2^46,47^. The transition from G2 to M phase requires a large increase in cyclin B/CDK1 activity^48,49^. The all-or-none checkpoints we documented in both G1 in and G2 are consistent with switch-like mechanisms that inhibit the intracellular pools of cyclin/CDK complexes below their respective thresholds necessary for cell cycle transitions. These mechanisms involve molecular entities such as the CDK inhibitor p21 and Wee1-mediated CDK inactivation^50–52^. Without the corresponding cyclin-CDK complex activities essential to initiating DNA replication or mitosis, cell cycle progression through the G1/S and S/G2 transitions is blocked rather than simply slowed.

In contrast, the graded slowdown kinetics we documented in S phase is consistent with the intra-S checkpoint mediated inhibition of late-origin firing. Complete genome duplication is normally achieved by establishing replication forks at thousands of individual DNA replication origins, but some origins fire at the beginning of S phase whereas others fire later. When DNA damage is detected (typically by stalled replication forks), nearby “dormant” origins that have not yet fired initiate replication to promote replication completion in that local region. At the same time, checkpoint pathways block new origin firing in other regions of the genome corresponding to late-replicating domains^53,54^. Thus, DNA synthesis continues at forks emanating from the early-fired origins, but the overall rate of S phase progression is slowed^55,56^. We postulate that this graded checkpoint response is protective to reduce the number of stalled replication forks that could collapse, snowballing the DNA damage^57^. Therefore, S phase actively counteracts the all-or-none mechanisms to avoid a prolonged stall with partially-replicated DNA, a situation that could lead to disastrous genome instability.

The location of the proposed commitment point can affect the apparent checkpoint stringency in a particular cell cycle phase. While both G1 and G2 utilize all-or-none checkpoint kinetics, the G2 checkpoint is more stringent than G1 in the sense that the commitment point occurs very close (within 0.5 hr) to the G2/M border. This temporal location of the G2 commitment point is consistent with previous observations and may be concurrent with the onset of antephase described previously using time-lapse analysis^58–62^. In contrast, the G1 commitment point is located ~3-4 hrs before the G1/S border in both RPE and U2OS, meaning that DNA damage within the last 3 hrs of G1 usually fails to delay S phase onset. The earlier commitment point in G1 is consistent with previous observations that the G1 checkpoint fails to prevent S phase entry at early times after irradiation^63^. This longer damage-insensitive period in G1 versus G2 may be explained by differences in their biochemical pathways. In G1, the DNA damage response (DDR) involves proteolysis of cyclin D and transcriptional changes such as p53-dependent p21 induction^64–66^, whereas DDR during G2 inactivates Cdc25, the rate-limiting step for CDK1 activation, through a faster series of phosphorylation and dephosphorylation reactions^67^. From an evolutionary standpoint, we speculate the earlier G1 commitment point from the G1/S transition might be advantageous in delaying DSB repair by the more error-prone non-homologous end joining (NHEJ) in G1 until S or G2, when the more error-free homologous recombination repair option is available^68–70^.

It is important to note that the commitment point that we describe is similar, but not identical, to the concept of a G1 restriction point, a temporal point at which cells in G1 are no longer sensitive to growth factor withdrawal for commitment to S phase entry^71^. Although mediated through different upstream mechanisms, the downstream consequences of serum starvation and DDR both converge on CDK2/4 inactivation^72^. Interestingly, the temporal location of the restriction point is comparable to the location of our G1 commitment point mapped 2-3 hours before DNA replication^73^, suggesting that different types of cell cycle checkpoints in G1 may act on a similar timescale due to their converging pathways. However, the DNA damage commitment point we identified is estimated to be earlier than the timing of the point-of-no-return (~0.7 hr before G1/S), which is characterized by irreversible APC inactivation^20^. However, we do not exclude the possibility that this discrepancy in timing might be explained by the use of different cell lines or methods to measure S phase onset (CDK2 activity versus PCNA foci formation). It is also possible that the location of the point-of-no-return may be stress type-specific, since a lower percentage of cells exit the cell cycle in response to DNA damage compared to oxidative and osmotic stresses^20^.

Previous studies that examine cell tolerance to DSBs, or radiosensitivity, measured by survival rate after irradiation, also exhibit drastic variation across different cell cycle phases. Specifically, G2 and M phase cells show the most sensitivity to radiation, followed by G1 cells, with S phase cells being the most resistant to damage^15^. Our results using a radiomimetic drug reveal a similar trend in checkpoint stringencies. The G2 checkpoint is the most stringent whereas the S phase checkpoint is the least stringent. Therefore, the stringencies of DNA damage checkpoints in each phase seem to be optimized to allow for maximal cell cycle progression while preventing the lethal consequences of DSBs. Consistent with previous cell cycle analyses in fixed cells^74,75^, S phase progression was minimally affected by damage except for extremely high NCS concentrations (Figure 3A-B). Even under the highest levels of damage, cells progressed through S phase and were rarely found to undergo permanent arrest within S phase. The fact that S phase shows the least stringent checkpoint yet is the most resistant to irradiation in terms of viability seems contradictory, since a robust DNA damage checkpoint would be expected to offer the strongest protection from irradiation. However, we found that cells damaged in S phase were arrested in the subsequent G2 phase by comparable or even higher probabilities than cells damaged directly during G2-M (**Figure S6**). This observation suggests that cells damaged in S phase carry incompletely repaired DSBs to G2 and then trigger a robust G2 checkpoint response that protects their viability. Therefore, therapeutically, it might be more effective to target S phase tumor cells when combined with checkpoint inhibitors of G2 rather than inhibitors of S phase progression.

Our work also demonstrates the advantage of using quantitative live-imaging approach to study cell cycle dynamics. Live-cell imaging measures DNA damage checkpoint dynamics under unperturbed conditions without artificial manipulations such as cell synchronization, which perturbs the cell cycle and can introduce unwanted DNA damage. Furthermore, the PCNA-based approach also provides an accurate methodology for locating cell cycle transitions and commitment points^12,21,76^. This method provides an advantage over the widely-used fluorescence ubiquitin cell cycle indicator (FUCCI) system^77,78,76,79^, a quantitative ubiquitination-based fluorescence reporter that uses tagged Cdt1 and geminin fragments to report the G1 and S-G2-M phases in real time. Whereas the FUCCI system is only capable of segmenting the cell cycle into 2 phases—G1 and S-G2-M^77–79^—and is not capable of sharply demarcating the G1/S transition^76^, the PCNA-based method provides a finer resolution demarking the boundaries at the G1/S and S/G2 borders^76^. The newer version of FUCCI (FUCCI4) was improved upon to visualize its S/G2 transition by adding a SLBP fragment and allowed for distinguishing G1, S, G2, and M with four distinct fluorescence colors in a snapshot without time lapse data. However, the identification of G1/S transition remained subjective and sensitive to thresholding^80^ with FUCCI4. This capability is crucial for precisely locating the proposed DNA damage commitment points relative to cell cycle phase boundaries. Another advantage of using PCNA as a marker of phase transition is its robustness to intensity fluctuation. In the FUCCI system, the demarcation of phase transitions relies on fluorescence intensity, which can vary under different imaging conditions, expression levels, and fluorophore degradation kinetics. In contrast, phase identification by the PCNA reporter relies on morphological changes due to protein localization that are insensitive to total fluorescence intensity as well as the rates of fluorophore maturation and degradation^33^. Thus, our results provide precise identification of the timing of cell cycle phase transitions after DNA damage.

Finally, our study reveals meaningful differences between an immortalized primary cell line and a transformed cell line derived from a human tumor. Despite qualitatively similar checkpoint dynamics observed within each phase between RPE and U2OS cells, we find quantitative differences in checkpoint stringencies—specifically, diminished checkpoint stringencies in the transformed U2OS cell lines across the cell cycle. This deficiency is most pronounced in G1 (Figure 5C, **Figure S6E-F**, highest NCS concentrations). U2OS cells are known to harbor an unstable G1 checkpoint due to p16 deficiency, truncated Wip1, and p53 heterozygosity^34,35,37^. Such diminished G1 checkpoint stringency is attributed to the earlier commitment point relative to the beginning of G1 (**Table S2**), halt duration upon checkpoint activation (**Figure S12B**), and diminished ability to permanently arrest cells (**Figure S12D**). Although both cell lines have a similar duration of the insensitive period after the commitment point, U2OS generally shows a significantly shorter total G1 duration than RPE. Thus, the G1 in U2OS cells has a relatively longer unchecked period compared to the G1 of RPE cells. In addition, our model estimates a better fit for the G1 checkpoint in U2OS with the graded slowdown, as opposed to the all-or-none kinetics seen in RPE cells (Table S2-3). It is possible that this difference in checkpoint kinetics may reflect a more general discrepancy between non-transformed and cancer cells. Graded slowdown kinetics may manifest as the “leakiness” in DNA damage checkpoint responses observed in transformed cells, which has been loosely attributed to defective checkpoint responses^81,82^. Thus, in light of cancer treatments that rely on differentially inducing DNA damage, it may be useful to clinically stratify tumors not only based on cell types, but also on the dynamics of DNA damage checkpoint responses.

## EXPERIMENTAL PROCEDURES

### Cell culture

U2OS cells were obtained from the laboratory of Dr. Yue Xiong and cultured in McCoy’s 5A medium supplemented with 10% fetal bovine serum (FBS) and penicillin/streptomycin (Gibco). hTERT retinal pigment epithelial cells (RPE) were obtained from the ATCC (ATCC^®^ CRL-4000^™^) and cultured in DMEM medium supplemented with 10% fetal calf serum and penicillin/streptomycin. When required, the medium was supplemented with selective antibiotics (2 μg/mL puromycin (Gibco)). When indicated, medium was replaced with fresh medium supplemented with neocarzinostatin (Sigma-Aldrich) during experiments. Clonogenic assay was performed using the CellTiter*-*Blue Cell Viability Assay (Promega). Cell cycle distributions were analyzed by DAPI staining and by incorporation of EdU using the Click-iT EdU kit (Invitrogen).

### Cell Line Construction

The pLenti-PGK-Puro-TK-NLS-mCherry-PCNA plasmid was subcloned from eGFP-PCNA (Gift from S. Angus) using the Gateway system (Life Technologies) following manufacturers protocols. The eGFP tag from the original plasmid was replaced with mCherry in the pENTR vector intermediate using standard methods. pLenti PGK Puro DEST (w529-2) was a gift from Eric Campeau (Addgene plasmid # 19068)^83^. The plasmid was stably expressed into U2OS or RPE cells by first transfecting the plasmid into 293T cells to generate replication-defective viral particles using standard protocols (TR-1003 EMD Millipore), which were used to stably infect the U2OS and RPE cell lines. The cells were maintained in selective media and hand-picked to generate a clonal population.

### Time-Lapse Microscopy

Prior to microscopy, cells were plated in poly-D-lysine coated glass-bottom plates (Cellvis) with FluoroBrite™ DMEM (Invitrogen) supplemented with FBS, 4 mM L-glutamine, and penicillin/streptomycin. Fluorescence images were acquired using a Nikon Ti Eclipse inverted microscope with a Nikon Plan Apochromat Lambda 40X objective with a numerical aperture of 0.95 using an Andor Zyla 4.2 sCMOS detector. In addition, we employed the Nikon Perfect Focus System (PFS) in order to maintain focus of live cells throughout the entire acquisition period. The microscope was surrounded by a custom enclosure (Okolabs) in order to maintain constant temperature (37°C) and atmosphere (5% CO_2_). The filter set used for mCherry was: 560/40 nm; 585 nm; 630/75 nm (excitation; beam splitter; emission filter) (Chroma). Images were acquired every 20 mins for U2OS cells and every 10 minutes for RPE cells in the mCherry channel. We acquired 2-by-2 stitched large image for RPE cell. NIS-Elements AR software was used for image acquisition and analysis.

### Image Analysis

Image analysis was performed using a customized CellProfiler pipeline that included speckle counting and object tracking^84^, followed by MATLAB (MathWorks) based custom written software to generate the PCNA foci trajectories in Figure 2E-F. Specifically, the CellProfiler pipeline utilized the PCNA-mCherry fluorescence channel to segment the nuclei, followed by speckle counting and object tracking to quantify PCNA-mCherry foci over time in each single nuclei. Because the success rate of the CellProfiler’s automatic cell tracking was unacceptably low (<20%), mostly due to the mobility of the cells, to obtain higher sample size, we manually tracked each cell and recorded the frame at which PCNA foci appeared (G1/S) or disappeared (S/G2) using ImageJ to produce all the transition curves.

### In silico mapping of cell cycle progression in individual cells

We quantified the cell cycle phase durations of our cell lines by imaging asynchronously dividing cells. During the entire life of each individual cell, we took four time point measurements: the time of cell birth (t_birth_), the onset of S phase (t_s_onset_), the end of S phase (t_s_end_), and the time of telophase (t_telophase_) (Figure 2A), which were manually identified from the PCNA-mCherry reporter. These four time points allowed for quantifying the durations of three major cell cycle phases: G1, S, and G2-M phases. For cells with exogenous DNA damage, prior to addition of NCS, we imaged freely cycling cells for at least 15 hours to establish the cell cycle phase duration before the DNA damage was induced.

### Immunofluorescence

Cells were fixed and permeabilized with -20°C methanol for 10 minutes, and stained overnight at 4°C with anti-phospho-H2AX Ser139 (JBW301, EMD Millipore 05-636). Primary antibodies were visualized using a secondary antibody conjugated to Alexa Fluor-488/-647 and imaged with appropriate filters. EdU incorporation and staining was performed using the Click-iT™ EdU kit (Invitrogen C10337).

## ACKNOWLEDGEMENTS

We thank Yue Xiong for helpful discussion, Samuel Wolff for guidance on experiments and microscopy operation, Katarzyna Kedziora for critical feedback, Chi Pham for editing the manuscript, and Po-Hao Huang for brainstorming ideas. This work was supported by National Institutes of Health research grants R00-GM102372 (JEP) and R01-GM102413 (JGC), National Institutes of Health training grants T32CA009156-40 (GDG), T32GM067553-12 (HXC), 1F30CA213876-01 (HXC), the North Carolina University Cancer Research Fund, and by a medical research grant from the W. M. Keck Foundation (JEP and JGC).

## AUTHOR CONTRIBUTIONS

HXC and GDG constructed the PCNA-mCherry reporter cell lines. HXC performed validation studies. HXC and JEP designed the experiments. HXC, CEP, AAP, and HYC performed live-cell imaging and experiments. HXC, CEP, AAP, and HYC conducted image analysis and cell tracking. HXC performed computational modeling and analysis. HXC wrote the manuscript with contributions from all authors.

## References

Nyberg, K. A., Michelson, R. J., Putnam, C. W. & Weinert, T. A. Toward Maintaining the Genome: DNA Damage and Replication Checkpoints. Annu. Rev. Genet. 36, 617–656 (2002).

Löbrich, M. & Jeggo, P. A. The impact of a negligent G2/M checkpoint on genomic instability and cancer induction. Nat. Rev. Cancer 7, 861–869 (2007).

Sogo, J. M., Lopes, M. & Foiani, M. Fork Reversal and ssDNA Accumulation at Stalled Replication Forks Owing to Checkpoint Defects. Science (80-.). 297, (2002).

Vilenchik, M. M. & Knudson, A. G. Endogenous DNA double-strand breaks: production, fidelity of repair, and induction of cancer. Proc. Natl. Acad. Sci. U. S. A. 100, 12871–6 (2003).

Haber, J. E. DNA recombination: the replication connection. Trends Biochem. Sci. 24, 271–5 (1999).

Bartek, J. & Lukas, J. DNA damage checkpoints: from initiation to recovery or adaptation. Curr. Opin. Cell Biol. 19, 238–245 (2007).

Wang, H., Zhang, X., Teng, L. & Legerski, R. J. DNA damage checkpoint recovery and cancer development. Exp. Cell Res. 334, 350–358 (2015).

Paulovich, A. G., Toczyski, D. P. & Hartwell, L. H. When checkpoints fail. Cell 88, 315–21 (1997).

Shaltiel, I. A. et al. The same, only different - DNA damage checkpoints and their reversal throughout the cell cycle. J. Cell Sci. 128, 607–20 (2015).

Finn, K., Lowndes, N. F. & Grenon, M. Eukaryotic DNA damage checkpoint activation in response to double-strand breaks. Cell. Mol. Life Sci. 69, 1447–73 (2012).

Arora, M., Moser, J., Phadke, H., Basha, A. A. & Spencer, S. L. Endogenous Replication Stress in Mother Cells Leads to Quiescence of Daughter Cells. Cell Rep. 19, 1351–1364 (2017).

Barr, A. R. et al. DNA damage during S-phase mediates the proliferation-quiescence decision in the subsequent G1 via p21 expression. Nat. Commun. 8, 14728 (2017).

Khanna, K. K. & Jackson, S. P. DNA double-strand breaks: signaling, repair and the cancer connection. Nat. Genet. 27, 247–54 (2001).

van Gent, D. C., Hoeijmakers, J. H. & Kanaar, R. Chromosomal stability and the DNA double-stranded break connection. Nat. Rev. Genet. 2, 196–206 (2001).

Pawlik, T. M. & Keyomarsi, K. Role of cell cycle in mediating sensitivity to radiotherapy. Int. J. Radiat. Oncol. Biol. Phys. 59, 928–42 (2004).

Elledge, S. J. Cell Cycle Checkpoints: Preventing an Identity Crisis. Science (80-.). 274, (1996).

Pomerening, J. R., Sontag, E. D. & Ferrell, J. E. Building a cell cycle oscillator: hysteresis and bistability in the activation of Cdc2. Nat. Cell Biol. 5, 346–51 (2003).

Novak, B., Tyson, J. J., Gyorffy, B. & Csikasz-Nagy, A. Irreversible cell-cycle transitions are due to systems-level feedback. Nat. Cell Biol. 9, 724–728 (2007).

L?pez-Avil?s, S., Kapuy, O., Nov?k, B. & Uhlmann, F. Irreversibility of mitotic exit is the consequence of systems-level feedback. Nature 459, 592–595 (2009).

Cappell, S. D., Chung, M., Jaimovich, A., Spencer, S. L. & Meyer, T. Irreversible APCCdh1 Inactivation Underlies the Point of No Return for Cell-Cycle Entry. Cell 166, 167–180 (2016).

Barr, A. R. et al. A Dynamical Framework for the All-or-None G1/S Transition. Cell Syst. 2, 27–37 (2016).

Branzei, D. & Foiani, M. Regulation of DNA repair throughout the cell cycle. Nat. Rev. Mol. Cell Biol. 9, 297–308 (2008).

Davis, D. M. & Purvis, J. E. Computational analysis of signaling patterns in single cells. Semin. Cell Dev. Biol. 37, 35–43 (2015).

Kisielewska, J., Lu, P. & Whitaker, M. GFP-PCNA as an S-phase marker in embryos during the first and subsequent cell cycles. Biol. Cell 97, 221–9 (2005).

Zölzer, F., Basu, O., Devi, P. U., Mohanty, S. P. & Streffer, C. Chromatin-bound PCNA as S-phase marker in mononuclear blood cells of patients with acute lymphoblastic leukaemia or multiple myeloma. Cell Prolif. 43, 579–83 (2010).

Sporbert, A., Gahl, A., Ankerhold, R., Leonhardt, H. & Cardoso, M. C. DNA Polymerase Clamp Shows Little Turnover at Established Replication Sites but Sequential De Novo Assembly at Adjacent Origin Clusters. Mol. Cell 10, 1355–1365 (2002).

Pomerening, J. R., Ubersax, J. A., Ferrell, J. E. & Jr. Rapid cycling and precocious termination of G1 phase in cells expressing CDK1AF. Mol. Biol. Cell 19, 3426–41 (2008).

Leonhardt, H. Dynamics of DNA Replication Factories in Living Cells. J. Cell Biol. 149, 271–280 (2000).

Essers, J. et al. Nuclear dynamics of PCNA in DNA replication and repair. Mol. Cell. Biol. 25, 9350–9 (2005).

Madsen, P. & Celis, J. E. S-phase patterns of cyclin (PCNA) antigen staining resemble topographical patterns of DNA synthesis. A role for cyclin in DNA replication? FEBS Lett. 193, 5–11 (1985).

Burgess, A., Lorca, T. & Castro, A. Quantitative live imaging of endogenous DNA replication in mammalian cells. PLoS One 7, e45726 (2012).

Trembecka-Lucas, D. O. & Dobrucki, J. W. A heterochromatin protein 1 (HP1) dimer and a proliferating cell nuclear antigen (PCNA) protein interact in vivo and are parts of a multiprotein complex involved in DNA replication and DNA repair. Cell Cycle 11, 2170–5 (2012).

Zerjatke, T. et al. Quantitative Cell Cycle Analysis Based on an Endogenous All-in-One Reporter for Cell Tracking and Classification. Cell Rep. 19, 1953–1966 (2017).

Diller, L. et al. p53 functions as a cell cycle control protein in osteosarcomas. Mol. Cell. Biol. 10, 5772–81 (1990).

Stott, F. J. et al. The alternative product from the human CDKN2A locus, p14(ARF), participates in a regulatory feedback loop with p53 and MDM2. EMBO J. 17, 5001–14 (1998).

Akan, P. et al. Comprehensive analysis of the genome transcriptome and proteome landscapes of three tumor cell lines. Genome Med. 4, 86 (2012).

Kleiblova, P. et al. Gain-of-function mutations of PPM1D/Wip1 impair the p53-dependent G1 checkpoint. J. Cell Biol. 201, 511–521 (2013).

Soltani, M., Vargas-Garcia, C. A., Antunes, D. & Singh, A. Intercellular Variability in Protein Levels from Stochastic Expression and Noisy Cell Cycle Processes. PLoS Comput. Biol. 12, e1004972 (2016).

Jullien, D., Vagnarelli, P., Earnshaw, W. C. & Adachi, Y. Kinetochore localisation of the DNA damage response component 53BP1 during mitosis. J. Cell Sci. 115, 71–9 (2002).

Cesare, A. J. Mitosis, double strand break repair, and telomeres: a view from the end: how telomeres and the DNA damage response cooperate during mitosis to maintain genome stability. Bioessays 36, 1054–61 (2014).

Orthwein, A. et al. Mitosis Inhibits DNA Double-Strand Break Repair to Guard Against Telomere Fusions. Science (80-.). 344, 189–193 (2014).

Heijink, A. M., Krajewska, M. & van Vugt, M. A. T. M. The DNA damage response during mitosis. Mutat. Res. Mol. Mech. Mutagen. 750, 45–55 (2013).

Zirkle, R. E. & Bloom, W. Irradiation of parts of individual cells. Science 117, 487–93 (1953).

Rieder, C. L. & Cole, R. W. Entry into Mitosis in Vertebrate Somatic Cells Is Guarded by a Chromosome Damage Checkpoint That Reverses the Cell Cycle When Triggered during Early but Not Late Prophase. J. Cell Biol. 142, 1013–1022 (1998).

Mikhailov, A. et al. DNA damage during mitosis in human cells delays the metaphase/anaphase transition via the spindle-assembly checkpoint. Curr. Biol. 12, 1797–806 (2002).

Labib, K. How do Cdc7 and cyclin-dependent kinases trigger the initiation of chromosome replication in eukaryotic cells? Genes Dev. 24, 1208–1219 (2010).

Tanaka, S. & Araki, H. Helicase Activation and Establishment of Replication Forks at Chromosomal Origins of Replication. Cold Spring Harb. Perspect. Biol. 5, a010371–a010371 (2013).

Taylor, W. R. & Stark, G. R. Regulation of the G2/M transition by p53. Oncogene 20, 1803–1815 (2001).

Lindqvist, A., Rodríguez-Bravo, V. & Medema, R. H. The decision to enter mitosis: feedback and redundancy in the mitotic entry network. J. Cell Biol. 185, 193–202 (2009).

Donzelli, M. & Draetta, G. F. Regulating mammalian checkpoints through Cdc25 inactivation. EMBO Rep. 4, 671–677 (2003).

Reinhardt, H. C. & Yaffe, M. B. Kinases that control the cell cycle in response to DNA damage: Chk1, Chk2, and MK2. Curr. Opin. Cell Biol. 21, 245–55 (2009).

Nigg, E. A. Mitotic kinases as regulators of cell division and its checkpoints. Nat. Rev. Mol. Cell Biol. 2, 21–32 (2001).

Ge, X. Q. & Blow, J. J. Chk1 inhibits replication factory activation but allows dormant origin firing in existing factories. J. Cell Biol. 191, 1285–1297 (2010).

Yekezare, M., Gomez-Gonzalez, B. & Diffley, J. F. X. Controlling DNA replication origins in response to DNA damage - inhibit globally, activate locally. J. Cell Sci. 126, 1297–1306 (2013).

Paulovich, A. G. & Hartwell, L. H. A checkpoint regulates the rate of progression through S phase in S. cerevisiae in response to DNA damage. Cell 82, 841–7 (1995).

Tercero, J. A. & Diffley, J. F. X. Regulation of DNA replication fork progression through damaged DNA by the Mec1/Rad53 checkpoint. Nature 412, 553–557 (2001).

Cortez, D. Preventing replication fork collapse to maintain genome integrity. DNA Repair (Amst). 32, 149–57 (2015).

Müllers, E., Silva Cascales, H., Burdova, K., Macurek, L. & Lindqvist, A. Residual Cdk1/2 activity after DNA damage promotes senescence. Aging Cell 16, 575–584 (2017).

Krenning, L., Feringa, F. M., Shaltiel, I. A., van den Berg, J. & Medema, R. H. Transient Activation of p53 in G2 Phase Is Sufficient to Induce Senescence. Mol. Cell 55, 59–72 (2014).

Müllers, E., Silva Cascales, H., Jaiswal, H., Saurin, A. T. & Lindqvist, A. Nuclear translocation of Cyclin B1 marks the restriction point for terminal cell cycle exit in G2 phase. Cell Cycle 13, 2733–43 (2014).

Xu, N. et al. Akt/PKB suppresses DNA damage processing and checkpoint activation in late G2. J. Cell Biol. 190, 297–305 (2010).

Feringa, F. M. et al. Hypersensitivity to DNA damage in antephase as a safeguard for genome stability. Nat. Commun. 7, 12618 (2016).

Deckbar, D. et al. The Limitations of the G1-S Checkpoint. Cancer Res. 70, 4412–4421 (2010).

Macleod, K. F. et al. p53-dependent and independent expression of p21 during cell growth, differentiation, and DNA damage. Genes Dev. 9, 935–44 (1995).

Waldman, T., Kinzler, K. W. & Vogelstein, B. p21 Is Necessary for the p53-mediated G1 Arrest in Human Cancer Cells. Cancer Res. 55, (1995).

Agami, R. & Bernards, R. Distinct Initiation and Maintenance Mechanisms Cooperate to Induce G1 Cell Cycle Arrest in Response to DNA Damage. Cell 102, 55–66 (2000).

Donzelli, M. & Draetta, G. F. Regulating mammalian checkpoints through Cdc25 inactivation. EMBO Rep. 4, 671–7 (2003).

Shrivastav, M., De Haro, L. P. & Nickoloff, J. A. Regulation of DNA double-strand break repair pathway choice. Cell Res. 18, 134–147 (2008).

Lieber, M. R., Ma, Y., Pannicke, U. & Schwarz, K. Mechanism and regulation of human non-homologous DNA end-joining. Nat. Rev. Mol. Cell Biol. 4, 712–720 (2003).

Aylon, Y., Liefshitz, B. & Kupiec, M. The CDK regulates repair of double-strand breaks by homologous recombination during the cell cycle. EMBO J. 23, 4868–4875 (2004).

Pardee, A. B. A Restriction Point for Control of Normal Animal Cell Proliferation. Proc. Natl. Acad. Sci. 71, 1286–1290 (1974).

Blagosklonny, M. V. & Pardee, A. B. The Restriction Point of the Cell Cycle. Cell Cycle 1, 102–109 (2002).

Campisi, J., Medrano, E. E., Morreo, G. & Pardee, A. B. Restriction point control of cell growth by a labile protein: evidence for increased stability in transformed cells. Proc. Natl. Acad. Sci. U. S. A. 79, 436–40 (1982).

Nüsse, M. Cell cycle kinetics of irradiated synchronous and asynchronous tumor cells with DNA distribution analysis and BrdUrd-Hoechst 33258-technique. Cytometry 2, 70–79 (1981).

Agarwal, M. L. et al. A p53-dependent S-phase checkpoint helps to protect cells from DNA damage in response to starvation for pyrimidine nucleotides. Proc. Natl. Acad. Sci. U. S. A. 95, 14775–80 (1998).

Wilson, K. A., Elefanty, A. G., Stanley, E. G. & Gilbert, D. M. Spatio-temporal re-organization of replication foci accompanies replication domain consolidation during human pluripotent stem cell lineage specification. Cell Cycle 0 (2016). doi:10.1080/15384101.2016.1203492

Saitou, T. & Imamura, T. Quantitative imaging with Fucci and mathematics to uncover temporal dynamics of cell cycle progression. Dev. Growth Differ. 58, 6–15 (2016).

Dowling, M. R. et al. Stretched cell cycle model for proliferating lymphocytes. Proc. Natl. Acad. Sci. U. S. A. 111, 6377–82 (2014).

Sandler, O. et al. Lineage correlations of single cell division time as a probe of cell-cycle dynamics. Nature 519, 468–471 (2015).

Bajar, B. T. et al. Fluorescent indicators for simultaneous reporting of all four cell cycle phases. Nat. Methods 13, 993–996 (2016).

Kastan, M. B. et al. A mammalian cell cycle checkpoint pathway utilizing p53 and GADD45 is defective in ataxia-telangiectasia. Cell 71, 587–597 (1992).

Bunz, F. et al. Requirement for p53 and p21 to Sustain G2 Arrest After DNA Damage. Science (80-.). 282, (1998).

Campeau, E. et al. A Versatile Viral System for Expression and Depletion of Proteins in Mammalian Cells. PLoS One 4, e6529 (2009).

Carpenter, A. E. et al. CellProfiler: image analysis software for identifying and quantifying cell phenotypes. Genome Biol. 7, R100 (2006).

